# Characterization of nuclear genome size and variation in a freshwater snail model system featuring a recent whole-genome duplication

**DOI:** 10.1101/2025.01.23.634510

**Authors:** Maurine Neiman, Maria Pichler, Martin Haase, Dunja K. Lamatsch

## Abstract

Species are defined by unique nuclear genome characteristics like nucleotide composition, genomic structure, and genome size. These fundamental aspects of the nuclear genome can themselves be the object of natural selection. We here provide the first high-quality direct measurements of nuclear genome DNA content in a representative diverse sample of *Potamopyrgus antipodarum*, an Aotearoa New Zealand freshwater snail that is a textbook example of the maintenance of sexual reproduction in nature and is invasive worldwide. We used propidium-iodide-based flow cytometry to characterize nuclear DNA content, and its variation, in nearly 100 *P. antipodarum* from multiple populations representing both sexual and asexual individuals. We also estimated nuclear DNA content in multiple *P. estuarinus*, a closely related obligately sexual species. These data confirmed and extended earlier lines of evidence for polyploidy and variable genome size within asexual *P. antipodarum* and provided the first direct evidence for distinctly higher nuclear genome content in diploid (sexual) *P. antipodarum* relative to diploid sexual *P. estuarinus*. These data are consistent with genomic evidence for a recent whole-genome duplication (WGD) and subsequent and in-process rediploidization in *P. antipodarum*, setting the stage for use of *Potamopyrgus* as a model for WGD and its consequences.

## Introduction

The size, structure, and composition of the nuclear genome are together perhaps the defining trait of any given unique taxon, and each play central but distinct roles in shaping evolutionary trajectories and organismal ecology (e.g., Hanken and Wake 1993, Knight et al. 2005, Kang et al. 2015, Guignard et al. 2017, Meyer et al. 2021, Guo et al. 2024). While whole-genome sequencing efforts can definitively establish nucleotide composition, the physical size of the genome (the amount of DNA in an unreplicated somatic nucleus, here “nuclear DNA content”) and genomic structure (e.g., karyotype) cannot always be rigorously and accurately characterized by even high-quality nuclear genome sequence assemblies (Gregory 2005, Hjelman 2024). This issue is exacerbated by common genomic features like repetitive content and high levels of heterochromatism, both of which are often themselves associated with relatively large genomes (Hjelman et al. 2020). Because nuclear DNA content can differ between closely related taxa and even conspecifics (Blommaert 2020; e.g., Alvarez-Fuster et al. 1991, Neiman et al. 2011, Huang et al. 2014, Becher et al. 2021; also see https://www.genomesize.com/), one also needs to estimate nuclear DNA content in a representative sample of conspecifics to provide an accurate picture of nuclear genome size (including potential for variation) for any given species.

A simpler and older technology, flow cytometry (Gregory 2005, Hjelman 2024), when appropriate intercalating fluorochromes (e.g., propidium iodide, PI) and internal standardization are used (Bennett and Leitch 2005; e.g., Hjelman et al. 2019), is still the gold standard for providing accurate and direct estimates of nuclear genome DNA content (Blommaert 2020). Here, we use PI-based flow cytometry to provide the definitive estimates of nuclear genome DNA content across ploidy levels in an Aotearoa New Zealand snail species, *Potamopyrgus antipodarum*, that has risen to prominence because of notably wide within and across-population variation in reproductive mode and ploidy level (Lively 1987, Wallace 1992, Neiman et al. 2011, Paczesniak et al. 2013) and is invasive worldwide (Geist et al. 2022). We also use flow cytometry to provide nuclear DNA content estimates for an obligately sexual and diploid congener, *P. estuarinus*.

These new data on nuclear DNA content in *Potamopyrgus* are important in providing an accurate picture of a fundamentally important trait for an important model system. Our results also provide a distinct line for evidence in support of the hypothesis that *P. antipodarum* appears to have experienced a very recent whole-genome duplication (WGD) that might be in turn connected to its unusually frequent transitions to asexual reproduction (Logsdon et al. 2017, Fields et al. 2024, Jalinsky et al. in prep). With respect to the WGD, accurate characterization of genome size in *P. antipodarum* and a congener that reasonably represents the “ancestral” (pre-duplication) nuclear genomic DNA content (Fields et al. 2024) is a critical piece of understanding the causes and consequences of this recent dramatic change in genome structure.

### Methods: Flow Cytometry & Estimation of Nuclear DNA Content

We used flow cytometry to measure nuclear DNA content following the detergent-trypsin method and PI staining approach detailed in Stelzer et al. (2011), with minor modifications. Details regarding the specific standards used and when flow cytometry was conducted for each of our samples can be found in Supplemental Table 1. We quantified nuclear genome DNA content in 99 *P. antipodarum* and 10 *P. estuarinus*. The *P. antipodarum* we used were either collected from natural populations in New Zealand (N = 67, 11 sites) or an invasive population in Austria (N = 5, 1 site), sourced from laboratory-cultured lineages descended from single females originally collected from New Zealand lakes (N = 32, 8 lineages), or sampled from diverse and uncharacterized tanks of *P. antipodarum* kept in the Neiman lab (N = 4, 3 tanks) (Table 1).

We first followed standard procedures (e.g., Neiman et al. 2012) to determine whether a penis was present (male) or absent (female) and thus sex each snail. Next, we dissected away the head tissue of each snail and snap-froze the tissue in liquid nitrogen, while the body tissue was fixed for chromosome preparation in Carnoy fixative. The head was homogenized in a citrate buffer (3.4 mM Trisodium citrate dihydrate, Nonidet P40 at 0.1% v/v, 1.5 mM sperminetetrahydrochloride, 0.5 mM trishydroxymethylaminomethane, pH 7.6), and was then transferred to 750 μl stock solution in a 1 ml Kimble Dounce tissue grinder (Sigma Aldrich) and homogenized on ice with 20 strokes using the “tight” pestle of the homogenizer. As an internal standard of known genome size, we used the fruit fly *Drosophila melanogaster* (genotype ISO- 1, diploid nuclear DNA content: 0.350 pg, and genotype DGRP-208, diploid nuclear DNA content: 0.321 pg (Gregory 2024)) and/or self-collected female chicken (*Gallus gallus*) red blood cells (diploid nuclear DNA content = 2.50 pg; Vinogradov 1998). The same procedure used to prepare and run samples for snail flow cytometry was applied to head tissue dissected from each of 10 *D. melanogaster* females and to the 200µl of chicken red blood cells (“CRBC”).

A Pearson’s correlation analysis demonstrated that genome size estimates for the 55 samples for which we used both *D. melanogaster* and CRBC standards to estimate nuclear genome DNA content were virtually identical (*r* = 0.996, *p* < 0.001; Fig. 1). This high correspondence between CRBC and *D. melanogaster* is especially important in light of the fact that the small size of the fly heads translates into higher CVs for fluorescence (see below). We nevertheless prefer to use only *D. melanogaster* standards for our genome content estimates (N = 54 samples) because, unlike CRBC (2.5 pg), the nuclear DNA content of *D. melanogaster* (<< 1 pg) is much lower than for even diploid *P. antipodarum* or *P. estuarinus* (Figs. 1-3), facilitating the differentiation of snail vs. standard.

**Figure 1.**
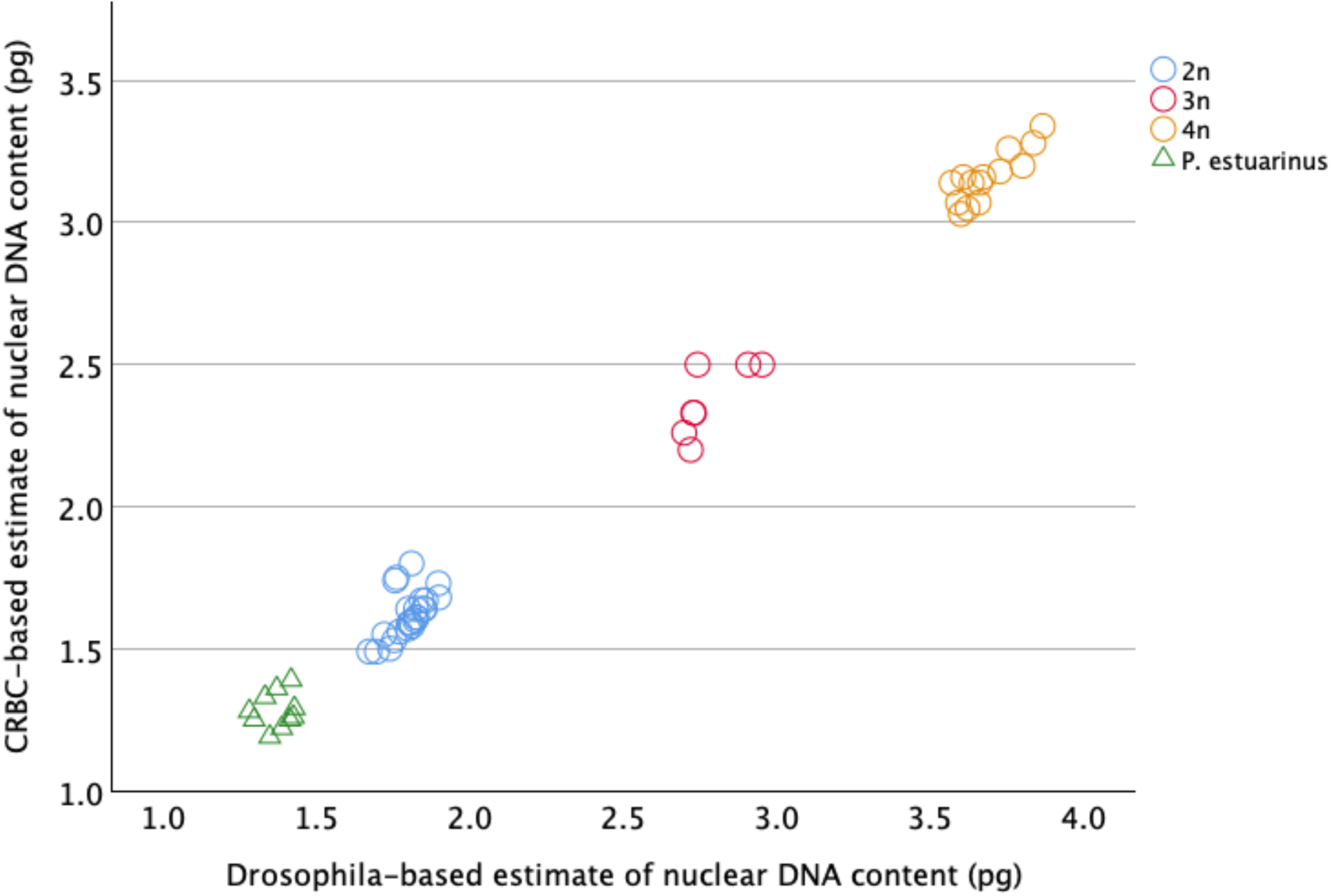
Scatterplot comparing the *Drosophila melanogaster*-based estimates of nuclear DNA content in *Potamopyrgus* relative to the chicken red blood cell-based measurements for the same samples. Open circles represent *P. antipodarum*, and the triangles represent *P. estuarinus*. Ploidy assignments reflect the position of each snail in the trimodal distribution shown in Fig. 2.

Large debris was removed by filtration through a 40-μm mesh nylon sieve before measurement. After addition of 100 μl of 0.021% Trypsin (dissolved in stock solution), the sample was incubated for exactly 10 min at 37°C. To stop the digestion, 75µl of 0.25% trypsin inhibitor was added (this solution also included 0.05% RNAse A), and the samples were incubated for another 10 min at 37°C. Finally, samples were stained with PI at a concentration of 50 μg/ml. Stained samples were kept overnight on ice in the dark. Flow cytometric analysis was performed on the next day on an Attune NxT^®^ acoustic focusing cytometer (Thermo Fisher) with an excitation wavelength of 561 nm and a custom-made 590–650 nm bandpass filter (yellow, YL-2) for detection of PI fluorescence. To exclude doublets (i.e., nuclei that pass the detector too close together, thus being recorded as a single “event”), we employed YL2-A *vs*. YL2-H gating. We first ran the homogenized and stained snail cells on their own to check quality (i.e., visual assessment of the variance around the snail fluorescence peak) without the standard. If the standard appeared to be of acceptable quality (i.e., this fluorescence peak is relatively narrow, suggesting low variance of fluorescence estimates), we then mixed the homogenized and stained cells from the snail and the standard and ran the sample again to provide a standard-calibrated estimate of nuclear genome content in the snail. We provide examples of these fluorescence peaks for diploid, triploid, and tetraploid *P. antipodarum* and for *P. estuarinus* in Figure 3.

**Figure 2.**
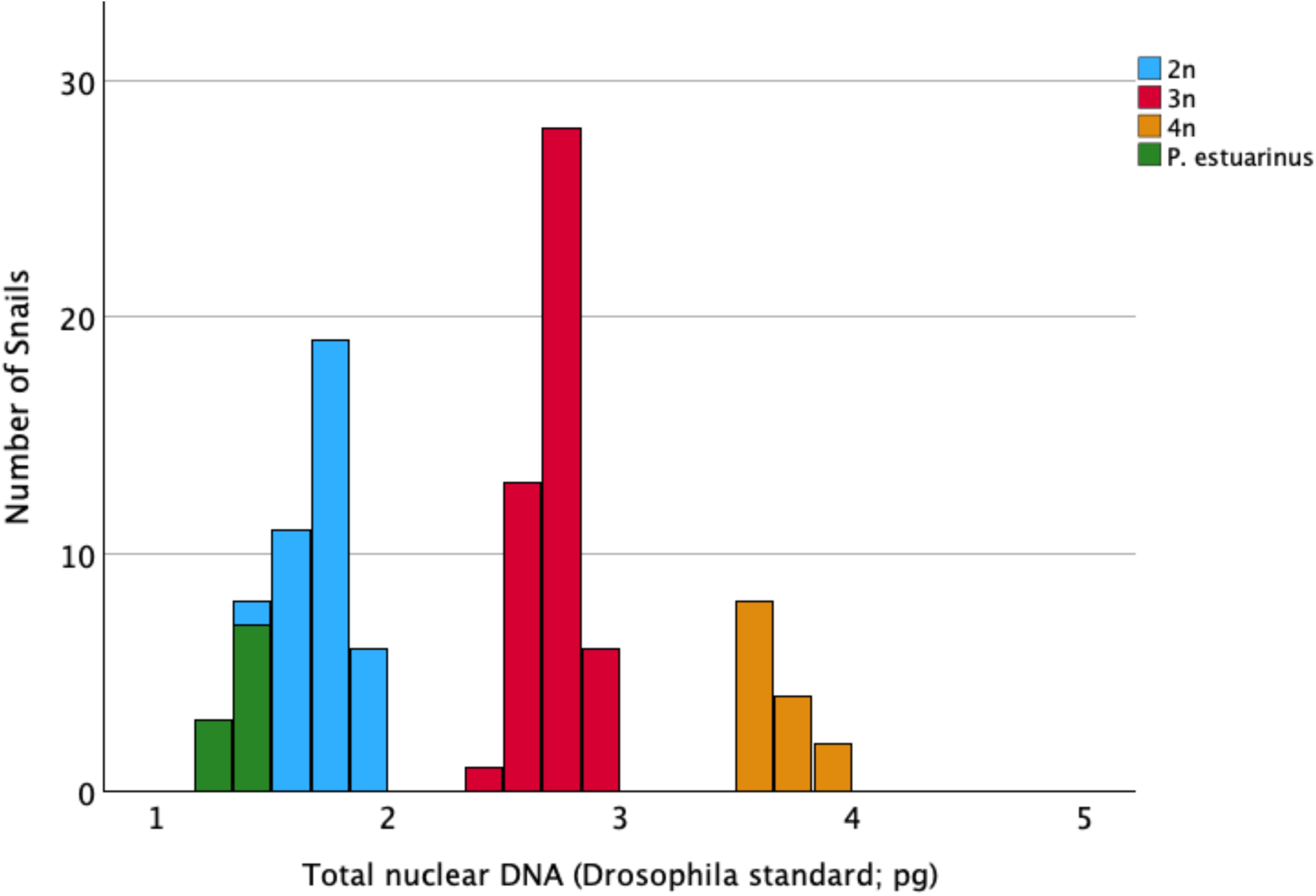
Histogram of genome size estimates across all 109 *Potamopyrgus*, color-coded according to species status (*P. antipodarum* vs. *P. estuarinus*) and ploidy (*P. antipodarum*), the latter assigned via position of data from each snail within the trimodal histogram for *P. antipodarum*. There is no evidence for ploidy variation within the diploid and obligately sexual *P. estuarinus*.

**Figure 3.**
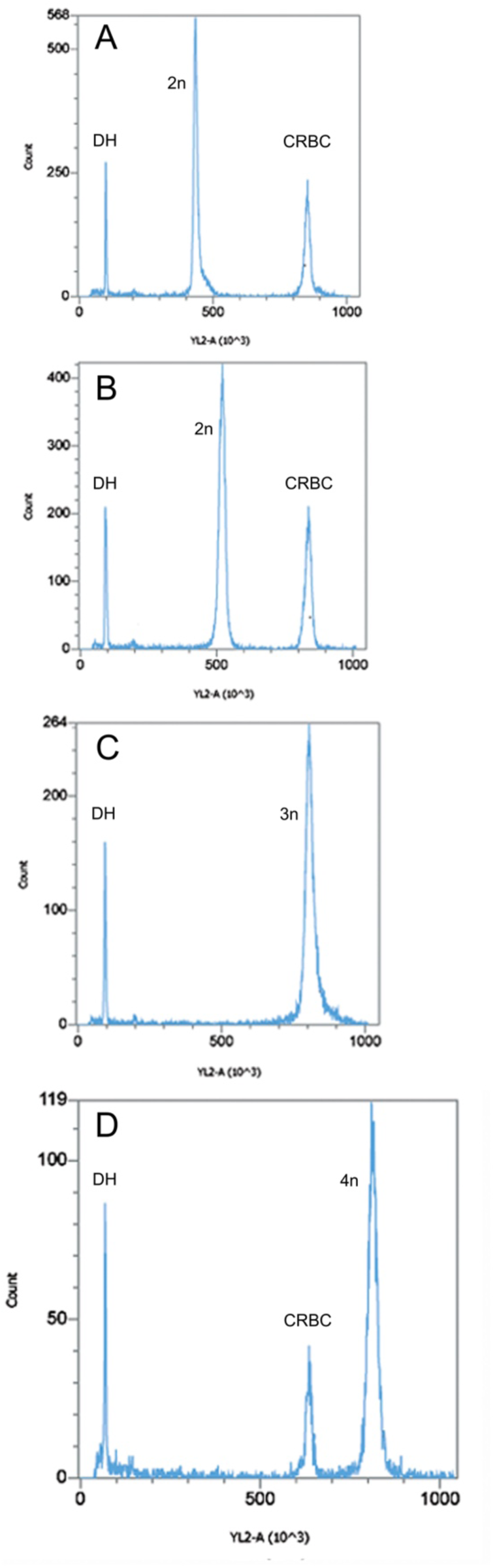
Examples of PI fluorescence peaks for (A) exemplar *P. estuarinus*, (B) diploid *P. antipodarum*, (C) triploid *P. antipodarum*, (D) tetraploid *P. antipodarum* (different scale, to accommodate higher nuclear DNA content). Each example also includes the standards spiked into that particular run, with “DH” peaks representing *Drosophila melanogaster* head standards and “CRBC” representing chicken red blood cell standards.

Coefficients of variance (CVs) of individual fluorescent peaks had a mean of 2.02% (SD ±0.77%) for snails, 1.48% (SD ± 0.18%) for CRBC, and 4.07% (SD ± 1.59%) for *D. melanogaster*. Conversion from picograms of DNA to base pairs were made following 1 pg = 978 Mbp (Gregory 2024). Following earlier flow cytometry-based studies in *P. antipodarum* (Neiman et al. 2011, Paczesniak et al. 2013), we visualized these nuclear genome DNA content data in the form of a histogram based on pg/nucleus. We thus assigned diploid, triploid, or tetraploid status for each *P. antipodarum* depending on where snail DNA content fit into the trimodal distribution of data (Fig. 2).

## Results & Discussion

We used propidium iodide-based flow cytometry to estimate nuclear genome DNA content for 99 *Potamopyrgus antipodarum*, a unique and prominent model system in ecology and evolution, and 10 *P. estuarinus*. This effort constitutes the first of which we are aware to use this gold-standard method for nuclear genome size quantification for a large sample of *P. antipodarum* from different populations, along with a congener often used for comparative studies. Such data are useful in themselves for any model system, but are of especially critical importance in providing an independent line of evidence suggesting that *P. antipodarum* has experienced a very recent whole-genome duplication (Logsdon et al. 2017; also see Fields et al. 2024).

As expected from earlier flow cytometry (Neiman et al. 2011, Paczesniak et al. 2013, Million et al. 2021), population genetic (Dybdahl and Lively 1995, Liu et al. 2012, Paczesniak et al. 2013), and cytogenetic data (Wallace 1992, McElroy et al. 2021), the distribution of nuclear DNA content within *P. antipodarum* was trimodal, corresponding to diploid, triploid, and tetraploid individuals (Fig. 2). Accordingly, we assigned each *P. antipodarum* individual to a ploidy level depending on its placement in one of the three nuclear DNA content peaks: diploid (mean nuclear DNA content = 1.710 pg ± 0.137 SD, N = 37), triploid (mean nuclear DNA content = 2.712 pg ± 0.103 SD, N = 48; 1.59x vs. diploids), or tetraploid (mean nuclear DNA content = 3.686 pg ±0.096 SD, N = 14; 2.16x vs. diploids and 1.36x vs. triploids) (Figs. 1, 2). The mean nuclear DNA content for the 10 *P. estuarinus* was 1.370 pg/nucleus (SD± 0.054), 80% of that of diploid *P. antipodarum* (Figs. 1, 2). Our PI-based estimate of nuclear genome DNA content in *P. estuarinus* is the first of which we are aware, but it is aligned with previous genomic sequencing-centered approaches to genome size estimation in this species (Fields et al. 2024; Supplemental Table 2). The distinctly higher nuclear DNA content of *P. antipodarum* relative to *P. estuarinus* provides yet another line of evidence pointing to a very recent whole-genome duplication in the former (Logsdon et al. 2017, Fields et al. 2024; Jalinsky et al. in prep).

In light of DAPI-based flow cytometry data that indicated wide variation in nuclear DNA content both across populations and within ploidies in *P. antipodarum* (Neiman et al. 2011) and similar variation within and among asexual triploid *P. antipodarum* from a single New Zealand lake (Million et al. 2021), we also compared nuclear DNA content across the 11 natural collections and 8 laboratory lineages in our *P. antipodarum* dataset. Similar to these earlier data, our PI-based estimates reveal variation within natural populations across ploidy levels (Te Anau, with 5 triploids and 1 tetraploid in our sample of 6 snails) as well as across snail sources within ploidy levels (Figs. 4, 5). We also see two cases of ploidy variation *within* laboratory lineages: there was 1 triploid snail in the otherwise diploid Selfe 18-1 lineage, and 1 triploid snail in the otherwise tetraploid Gunn-07 lineage.

**Figure 4.**
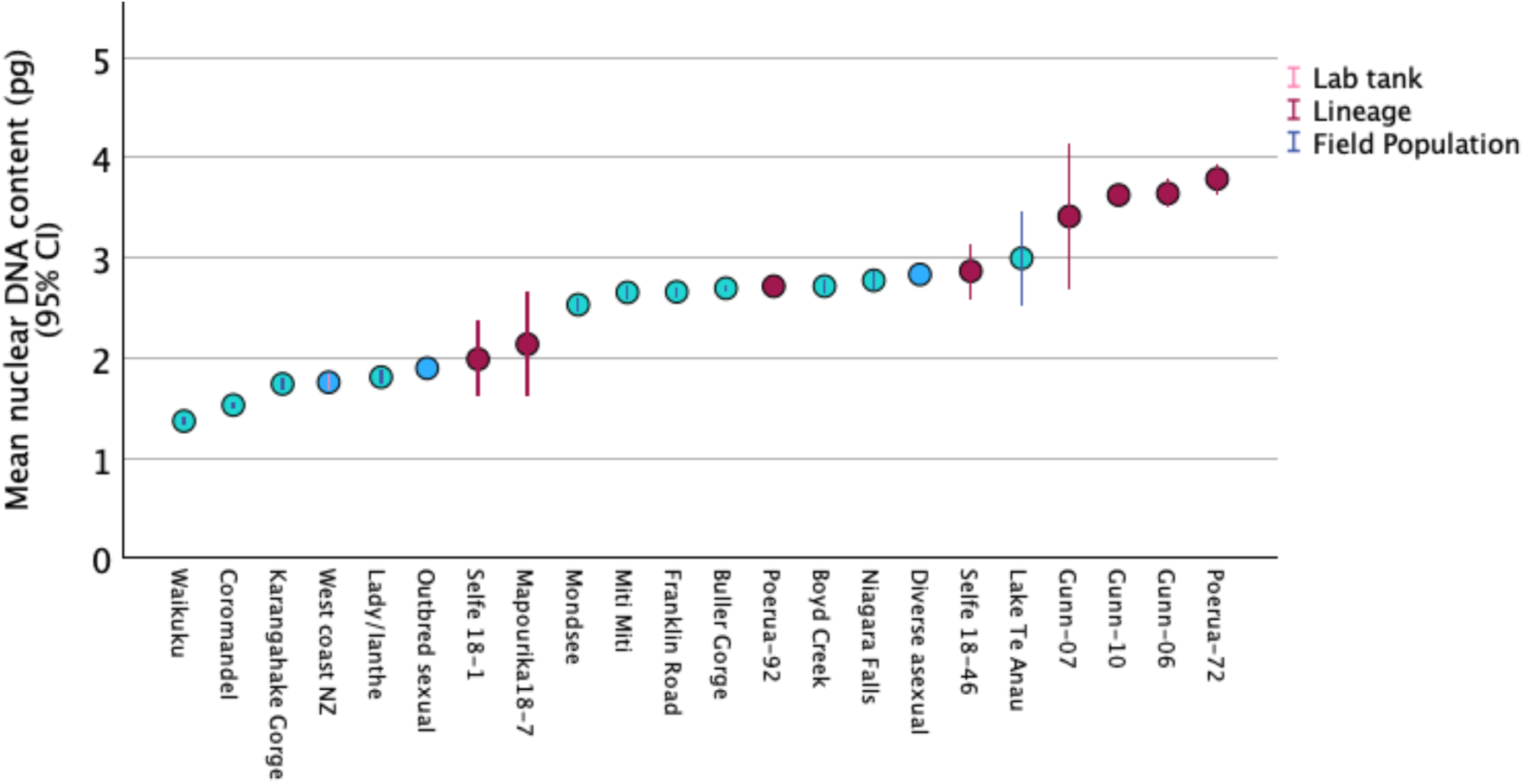
Comparisons across mean nuclear DNA content from snails collected from natural populations and laboratory lineages. The especially wide 95% CI around the means for Selfe 18- 1, Mapourika 18-7, Gunn-07, and Te Anau reflect ploidy polymorphism within the samples; all other samples were only represented by one ploidy level.

We used a bootstrapping approach as implemented within IBM SPSS v. 29 software to compare 95% confidence intervals across median nuclear genome content within ploidy levels and across snail sources (Supplemental Table 3). This analysis revealed that some *P. antipodarum* harbored significantly different nuclear DNA content even within ploidy levels, based on non-overlapping 95% CI around medians (Fig. 5). For example, Coromandel Peninsula 2n snails have significantly lower nuclear DNA content than all other 2n *P. antipodarum*, but still significantly higher nuclear DNA content than *P. estuarinus*, and 4n Poerua-72 snails have significantly higher nuclear DNA content than at least two of the three 4n lineages from lake Gunn. These outcomes are consistent with and extend earlier reports of such variation in *P. antipodarum* (Neiman et al. 2011, Million et al. 2021), which is turn potentially a consequence of variable chromosome number (Winterbourn 1970, Wallace 1992, McElroy et al. 2021) and/or variation in the extent of accumulation of an ∼18 kb tandemly repeated block of histone and ribosomal DNA genes (McElroy et al. 2021) or other repetitive elements (e.g., Blommaert et al. 2019). Definitive assessment of these non-mutually exclusive hypotheses will require cytogenetic analysis of individuals with the same assigned ploidy level but differing in nuclear genome DNA content.

**Figure 5.**
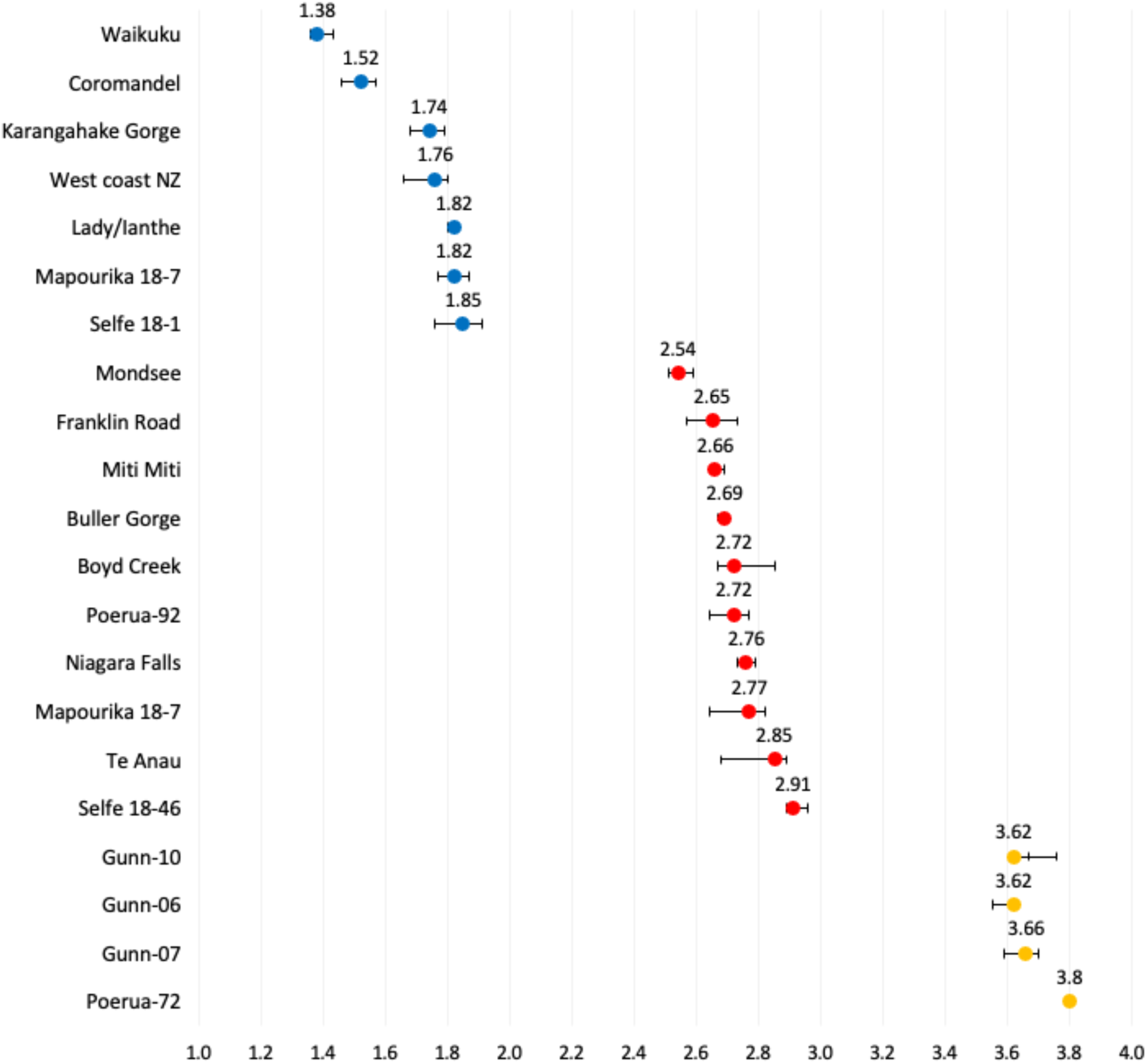
Comparison of median nuclear DNA content (pg; x axis) across source populations and lineages within ploidy levels. Samples are rank-ordered from top to bottom by median pg within ploidy level from low to high (Supplemental Table 3). Diploids are blue, triploids are red, and tetraploids are yellow. Only snail sources represented by more than one individual per ploidy level are included (see Table 1). “Waikuku” in the 2n panel is the *P. estuarinus* sample. Error bars represent 95% confidence intervals for median nuclear DNA content generated by 884-1000 bootstrap samples as executed within IBM SPSS (Supplemental Table 3).

The distinctly lower nuclear genome content of the snails from the Coromandel Peninsula relative to other presumably diploid and sexual *P. antipodarum* (Figs. 4, 5) prompted us to consider whether these individuals might instead represent a distinct species. We followed Haase (2008), who used careful morphological observation coupled with assessment of mitochondrial DNA sequence divergence relative to other *Potamopyrgus* taxa to demonstrate the presence of cryptic *Potamopyrgus* species in New Zealand in his 2008 revision of the radiation of tateid (at the time attributed to Hydrobiidae; see Wilke et al. 2013) gastropods in New Zealand. We dissected each of three Coromandel Peninsula males and females to characterize genital anatomies and assess oviparous vs. ovoviviparous status. We also sequenced a 503-bp fragment of mitochondrial *cytochrome b* [see Neiman and Lively (2004) for primers and PCR conditions] in an additional seven Coromandel Peninsula individuals to determine whether even phenotypically similar individuals might differ genetically.

Genital anatomy is diagnostic of species in *Potamopyrgus* (Haase 2008), and the fact that we did not detect any such anatomical differences with respect to *P. antipodarum* for the Coromandel Peninsula snails and that these snails were ovoviviparous indicate that the Coromandel Peninsula snails are *P. antipodarum*. This conclusion is bolstered by the mtDNA data: the two haplotypes detected in the 7 snails (identical in six individuals, differing at one position for one snail) were 98.21%-99.40% (Genbank accession PQ962506) and 98.21%-99.60% (PQ962507) identical to the first 30 hits of *P. antipodarum* in a BLAST search against the core nucleotide database of GenBank (21 January 2025; https://blast.ncbi.nlm.nih.gov/Blast.cgi). The closest hit between *P. antipodarum* and another *Potamopyrgus* species (*P. kaitunuparaoa*) was to 94.01% pairwise identity. Thus, neither genital anatomy, mode of egg production, nor mitochondrial sequence information indicated that the population from the Coromandel Peninsula, despite its distinctly smaller nuclear genome DNA content, represented a species other than *P. antipodarum*.

The discovery of a *P. antipodarum* population with smaller nuclear genome DNA content than what appears to be the majority of its otherwise similar diploid sexual counterparts sets the stage for follow-up work more directly evaluating the causes and consequences of the extensive genome size variation, both within and across ploidy levels, in this species. Especially productive next steps might include sequencing of nuclear markers that will allow us to verify or refute the tentative conclusion regarding the assignment of the Coromandel Peninsula snails as members of *P. antipodarum*, high-coverage genomic sequencing and cytogenetic characterization of Coromandel Peninsula snails to assess the extent of representation (or lack thereof) by repetitive elements (following McElroy et al. 2021), and crossing experiments between Coromandel Peninsula snails and diploid sexual *P. antipodarum* with the more “standard” higher nuclear DNA content followed by genomic and cytogenetic characterization of offspring.

## Summary, Conclusions, & Next Steps

We used PI-based quantification of nuclear genome DNA content in a diverse sample of *P. antipodarum* and multiple *P. estuarinus* to provide the first rigorous and comprehensive characterization of this fundamental genomic trait in a genus known around the world as a model system for the evolution of sex and host-parasite interactions and as a worldwide biological invader. These analyses validated and qualitatively extended previously established evidence for polyploidy and variable nuclear genome DNA content *within* ploidy levels in *P. antipodarum.* This work also provides a distinct line of support for a recent whole-genome duplication (WGD) in the lineage leading to *P. antipodarum* after its divergence from the shared common ancestor with *P. estuarinus.* Together, these data provide a concrete example of the importance of surveying a large and diverse sample of individuals with respect to accurately assessing intraspecific variation in nuclear genome DNA content and set the stage for the use of *Potamopyrgus* as a powerful system to explore questions regarding the causes and consequences of genome size variation and the aftermath of a recent WGD.

## Acknowledgments

Funding was provided by NSF MCB-1122176 and NSF DEB-1753851. Professor Manfred Schartl also contributed funds to support flow cytometry analyses. We thank the University of Iowa International Programs for International Travel Awards that supported travel for data collection and analysis in 2019 and 2021, and the University of Innsbruck for a guest professorship to Maurine Neiman in 2019. Some of the collections were made in the frame of the Research Training Group 2010 RESPONSE funded by the German Science Foundation. We thank Gerlien Verhaegen, Lisa Männer, Michael Winterbourn, John Logsdon, Mary Morgan-Richards, Curt Lively, Tim Linksvayer, Bennett Brown, JJ Neiman-Brown, Asheley Landrum, Katelyn Larkin, Laura Bankers, Kyle McElroy, and Jeremy Richardson for snail collections. We thank Marissa Roseman, Ben Ripperger, Mohammed Farooqi, and Silke Fregin for snail maintenance.

## References Cited

Alvarez-Fuster A, Juan C, Petitpierre E (1991) Genome size in *Tribolium* flour-beetles: inter- and intraspecific variation. Genetical Research 58:1–5.

Becher H, Powell RF, Brown MR, Metherell C, Pellicer J, Leitch IJ, Twyford AD (2021) The nature of intraspecific and interspecific genome size variation in taxonomically complex eyebrights. Annals of Botany 128:639–51.

Bennett MD, Leitch IJ (2005) Genome size evolution in plants. In: The Evolution of the Genome, p. 89-162. Elsevier Academic Press.

Blommaert J (2020) Genome size evolution: towards new model systems for old questions. Proceedings of the Royal Society B 1933 :20201441.

Blommaert J, Riss S, Hecox-Lea B, Mark Welch DB, Stelzer C-P (2019) Small, but surprisingly repetitive genomes: transposon expansion and not polyploidy has driven a doubling in genome size in a metazoan species complex. BMC Genomics 20:466.

Dybdahl MF, Lively CM (1995) Diverse, endemic and polyphyletic clones in mixed populations of a freshwater snail (*Potamopyrgus antipodarum*). Journal of Evolutionary Biology 8:385–98.

Fields PD, Jalinsky JR, Bankers L, Elroy KE, Sharbrough J, Higgins C, Morgan-Richards M, Boore JL, Neiman M, Logsdon JM Jr (2023) Genome evolution and introgression in the New Zealand mud snails *Potamopyrgus estuarinus* and *Potamopyrgus kaitunuparaoa*. Genome Biology and Evolution 16:evae091.

Geist JA, Mancuso JL, Morin MM, Bommarito KP, Bovee EN, Wendell D, Burroughs B, Luttenton MR, Strayer DL, Tiegs SD (2022) The New Zealand mud snail (*Potamopyrgus antipodarum*): autecology and management of a global invader. Biological Invasions 24:905–938.

Gregory TR (2005) In: The Evolution of the Genome, p. 3-87. Elsevier Academic Press.

Gregory TR (2024) Animal Genome Size Database. http://www.genomesize.com.

Guignard MS, Leitch AR, Acquisti C, Eizaguirre C, Elser JJ, Hessen DO, Jeyasingh PD, Neiman M, Richardson AE, Soltis PS, Soltis DE (2017) Impacts of nitrogen and phosphorus: from genomes to natural ecosystems and agriculture. Frontiers in Ecology and Evolution 5:70.

Guo K, Pyšek P, van Kleunen M, Kinlock NL, Lučanová M, Leitch IJ, Pierce S, Dawson W, Essl F, Kreft H, Lenzner B (2024) Plant invasion and naturalization are influenced by genome size, ecology and economic use globally. Nature Communications 15:1330.

Haase M (2008) The radiation of hydrobiid gastropods in New Zealand: a revision including the description of new species based on morphology and mtDNA sequence information. Systematics and Biodiversity 6:99–159.

Hanken J, Wake DB (1993) Miniaturization of body size: organismal consequences and evolutionary significance. Annual Review of Ecology and Systematics 1:501–19.

Hjelmen CE (2024) Genome size and chromosome number are critical metrics for accurate genome assembly assessment in Eukaryota. Genetics 227:iyae099.

Hjelmen CE, Blackmon H, Holmes VR, Burrus CG, Johnston JS (2019) Genome size evolution differs between *Drosophila* subgenera with striking differences in male and female genome size in Sophophora. G3: Genes, Genomes, Genetics 9:3167–79.

Huang W, Massouras A, Inoue Y, Peiffer J, Ràmia M, Tarone AM, Turlapati L, Zichner T, Zhu D, Lyman RF, Magwire MM (2014) Natural variation in genome architecture among 205 Drosophila melanogaster Genetic Reference Panel lines. Genome Research 24:1193–208.

Kang M, Wang J, Huang H (2015) Nitrogen limitation as a driver of genome size evolution in a group of karst plants. Scientific Reports 5:11636.

Knight CA, Molinari NA, Petrov DA (2005) The large genome constraint hypothesis: evolution, ecology and phenotype. Annals of Botany 95:177–90.

Lively CM (1987) Evidence from a New Zealand snail for the maintenance of sex by parasitism. Nature 328:519–21.

Logsdon J, Neiman M, Boore J, Sharbrough J, Bankers L, McElroy K, Jalinsky J, Fields P, Wilton P (2017) A very recent whole genome duplication in *Potamopyrgus antipodarum* predates multiple origins of asexuality & associated polyploidy. PeerJ Preprints: 2017 Jun 24.

Liu HP, Hershler R, Marn J, Worsfold TM (2012) Microsatellite evidence for tetraploidy in invasive populations of the New Zealand mudsnail, *Potamopyrgus antipodarum* (Gray, 1843). Journal of Molluscan Studies 78:227–30.

McElroy KE, Müller S, Lamatsch DK, Bankers L, Fields PD, Jalinsky JR, Sharbrough J, Boore JL, Logsdon Jr JM, Neiman M (2021) Asexuality associated with marked genomic expansion of tandemly repeated rRNA and histone genes. Molecular Biology and Evolution 38:3581–92.

Meyer A, Schloissnig S, Franchini P, Du K, Woltering JM, Irisarri I, Wong WY, Nowoshilow S, Kneitz S, Kawaguchi A, Fabrizius A (2021) Giant lungfish genome elucidates the conquest of land by vertebrates. Nature 590:284–9.

Million KM, Bhattacharya A, Dinges ZM, Montgomery S, Smith E, Lively CM (2021) DNA content variation and SNP diversity within a single population of asexual snails. Journal of Heredity 112:58–66.

Neiman M, Lively CM (2004) Pleistocene glaciation is implicated in the phylogeographical structure of *Potamopyrgus antipodarum*, a New Zealand snail. Molecular Ecology 13:3085–98.

Neiman M, Larkin K, Thompson AR, Wilton P (2012) Male offspring production by asexual *Potamopyrgus antipodarum*, a New Zealand snail. Heredity 109:57–62.

Neiman M, Paczesniak D, Soper DM, Baldwin AT, Hehman G (2011) Wide variation in ploidy level and genome size in a New Zealand freshwater snail with coexisting sexual and asexual lineages. Evolution 65:3202–16.

Paczesniak D, Jokela J, Larkin K, Neiman M (2013) Discordance between nuclear and mitochondrial genomes in sexual and asexual lineages of the freshwater snail *Potamopyrgus antipodarum*. Molecular Ecology 22:4695–710.

Stelzer C-P, Riss S, Stadler P (2011) Genome size evolution at the speciation level: The cryptic species complex *Brachionus plicatilis* (Rotifera). BMC Evolutionary Biology 11:90.

Vinogradov AE (1998) Genome size and GC-percent in vertebrates as determined by flow cytometry: the triangular relationship. Cytometry 31:100–109.

Wallace C (1992) Parthenogenesis, sex and chromosomes in *Potamopyrgus*. Journal of Molluscan Studies 58:93–107.

Wilke T, Haase M, Hershler R, Liu H-P, Misof B, Ponder W (2013) Pushing short DNA fragments to the limit: Phylogenetic relationships of ‘hydrobioid’ gastropods (Caenogastropoda: Rissooidea). Molecular Phylogenetics and Evolution 66:715–36.

